# Membrane-associated α-tubulin is less acetylated in postmortem prefrontal cortex from depressed subjects relative to controls: cytoskeletal dynamics, HDAC6 and depression

**DOI:** 10.1101/2020.01.22.915991

**Authors:** Harinder Singh, Justyna Chmura, Runa Bhaumik, Ghanshyam N. Pandey, Mark M. Rasenick

## Abstract

Cytoskeletal proteins and post-translational modifications play a role in mood disorders. Post-translational modifications of tubulin also alter microtubule dynamics. Furthermore, tubulin interacts closely with Gα_s_, the G-protein responsible for activation of adenylyl cyclase. Postmortem tissue derived from depressed suicide brain showed increased Gα_s_ in lipid-raft domains compared to normal subjects. Gα_s_, when ensconced in lipid-rafts, couples less effectively with adenylyl cyclase to produce cAMP and this is reversed by antidepressant treatment. A recent *in-vitro* study demonstrated that tubulin anchors Gα_s_ to lipid-rafts and that increased tubulin acetylation (due to HDAC-6 inhibition) and antidepressant treatment decreased the proportion of Gα_s_ complexed with tubulin. This suggested that deacetylated-tubulin might be more prevalent in depression. This study, examined tubulin acetylation in whole tissue homogenate, plasma-membrane and lipid-raft membrane domains in tissue from normal control (NC) subjects, depressed suicides and depressed non-suicides. While tissue homogenate showed no changes in 〈-tubulin/tubulin acetylation between control, depressed suicides and depressed non-suicides, plasma-membrane associated tubulin showed significant decreases in acetylation in depressed suicides and depressed non-suicides compared to controls. No change was seen in expression of the enzymes responsible for tubulin acetylation or deacetylation. These data suggest that during depression, membrane localized tubulin maintains a lower acetylation state, permitting increased sequestration of Gα_s_ in lipid-raft domains, where it is less likely to couple to adenylyl cyclase for cAMP production. Thus, membrane tubulin may play a role in mood disorders which could be exploited for diagnosis and treatment.

**Significance Statement:** There is little understanding about the molecular mechanisms involved in the development of depression and in severe cases, suicide. Evidence for the role of microtubule modifications in progression of depressive disorders is emerging. These postmortem data provide strong evidence for membrane tubulin modification leading to reduced efficacy of the G protein, Gsα, in depression. This study reveals a direct link between decreased tubulin acetylation in human depression and the increased localization of Gα_s_ in lipid-raft domains responsible for attenuated cAMP signaling. The evidence presented here suggest a novel diagnostic and therapeutic locus for depression.

## Introduction

Hallmarks of Major depressive disorder (MDD) include persistent sad mood, anhedonia, changes in appetite, disturbed sleep, feelings of worthlessness, hopelessness and suicidal thoughts. While various antidepressant drug therapies are available, the biological underpinnings of their action as well as the molecular events leading to depression remain uncertain. Numerous suggestions about the biology of depression exist, and epigenetics (histone deacetylases-HDACs) and HDAC inhibitors as novel antidepressants are a recent addition to this list (Tsankova et al., 2007; Covington et al., 2009; Gundersen and Blendy, 2009). The majority of the presently available antidepressants have among their actions, prevention of monoamine uptake or degradation and one consistent effect of antidepressant treatment has been a persistent increase in cAMP and an upregulation of the cAMP generating system (Nibuya et al., 1996; Malberg et al., 2000; Donati and Rasenick, 2003). Furthermore, PET studies from depressed subjects showed global decreases in brain cAMP and antidepressant drugs restored cAMP levels (Fujita et al., 2007; Fujita et al., 2017). We have suggested that antidepressants achieve this by a gradual removal of Gα_s_ from lipid rafts and increasing association of that molecule with adenylyl cyclase Zhang and Rasenick, 2010; Czysz et.al 2015). Consistent with this postmortem samples from depressed human subjects reveal increased Gα_s_ (Donati et al., 2008). Gα_s_ is the only heterotrimeric G protein undergoing translocation out of lipid-rafts in response to antidepressant treatment ((Toki et al., 1999; Donati and Rasenick, 2005). Interestingly, antidepressant drugs have been shown to concentrate in lipid raft domains (Eisensamer et al., 2005; Erb et al., 2016). Together, these studies suggest that the lipid environment of Gα_s_ may play an important role in its localization and function, and that chronic antidepressant treatment alters the membrane localization of Gα_s_, resulting in augmented coupling to adenylyl cyclase (Allen et al., 2009; Zhang and Rasenick, 2010).

There is evidence for a role of cytoskeletal (microtubules) alterations in the pathology of several neuropsychiatric diseases Perez et.al, 2009(Brown et al., 2013; Wong et al., 2013; Scifo et al., 2017). These disorders are associated with structural changes in brain including synaptic pruning defects and spine and dendrite atrophy (Glausier and Lewis, 2013). The development of depression is associated with exposure to triggering environmental factors such as chronic stress (Pittenger and Duman, 2008; Lin and Koleske, 2010; Schmitt et al., 2014) (McEwen et al., 2017). Most importantly, post-translational modifications such as acetylation of tubulin help to maintain cytoskeletal stability (Idriss, 2000; Westermann and Weber, 2003).

Lipid-raft domains are also associated with cytoskeletal elements such as microtubules. Tubulin is comprised of an αβ dimer, and these dimers are localized in membranes, and enriched in lipid-rafts. Upon activation, Gα_s_ is released form the membrane, where it binds tubulin, activates tubulin GTPase and increases microtubule dynamics (Roychowdhury and Rasenick, 1994; Dave et al., 2011; Sarma et al., 2015). These findings suggest that tubulin may act as an anchor for Gα_s_ within the lipid-raft domains. A recent *in vitro* study (Singh et al., 2018) shows that treatment with antidepressants reduces the extent to which Gα_s_ is complexed with tubulin.

The enzymes responsible for the regulation of acetylation status of α-tubulin are histone deacetylase-6 (HDAC-6; deacetylating) and alpha-tubulin acetyl transferase-1 (ATAT-1: acetylating). There is emerging evidence for the role of HDAC in neuropsychiatric disorders, including MDD (Guidotti A et al., 2011; Tsankova N et al., 2007; Hobara T et al., 2010). Altered levels of HDAC 2, 4, 5, 6, 8 mRNA have been observed in blood cells and postmortem brain from mood disorder subjects (Guidoti A et al., 2011; Hubbert C et al., 2002). HDAC-6, localized in cytosol, deacyates α-tubulin (Hubbert C et al., 2002; Verdel A et al., 2000). Peripheral white blood cells derived from MDD subjects showed altered HDAC6 mRNA levels (Hobara T et al., 2010).

The current study compares the acetylation status of α-tubulin from postmortem human brain of depressed subjects and controls without known psychiatric histories. Prefrontal cortex (PFC) tissue showed comparable tubulin acetylation in homogenates, but strikingly decreased acetylation in membranes prepared from depressed suicides and depressed non-suicides. These data correspond well with a previous study showing increased Gα_s_ levels in lipid rafts, since acetylation of tubulin decreases its ability to bind Gα_s_ and anchor it to lipid rafts, resulting in less Gα_s_ available for adenylyl cyclase activation in the depressed brain. These findings also parallel those of Gα_s_ translocation from lipid-rafts by HDAC6 inhibitors (Singh et al., 2018). The data presented here and previous studies in model systems suggest that Gα_s_ anchoring to lipid rafts is involved in both depression and therapies for that disease through modulation of the cAMP-generating system. These findings suggest a direct role of HDAC6 in maintaining acetylation status of α-tubulin, stabilizing/destabilizing microtubules during normal and depressive states. The data also suggest that tubulin acetylation may be relevant to depression and its treatment.

## Materials and Methods

### Human Subject Information

Tissue used in this study was from Brodmann area 9 obtained from the right hemisphere of depressed suicide subjects (*n* = 15), depressed non-suicide subjects (*n* = 12) and normal control subjects (*n* = 15). Subject demographics are described in Table 1.. Brain tissues were obtained from the Maryland Brain Collection at the Maryland Psychiatric Research Center (Baltimore, MD). Tissues were collected only after a family member gave informed consent. All procedures were approved by the University of Maryland Institutional Review Board (IRB) and by the University of Illinois IRB.

**Table 1:** Demographic characteristics of suicide and control subjects

| Group No. | Group | Diagnosis | Age | Race | Gender | PMI (hr) | Brain pH | Cause of Death | Drug Toxicity | Antidepressants Yes/No |
| --- | --- | --- | --- | --- | --- | --- | --- | --- | --- | --- |
| 1 | CONTROL | NORMAL | 37 | Black | Male | 5 | 7.1 | ASCVD | None | No |
| 2 | CONTROL | NORMAL | 31 | Black | Male | 8 | 7.2 | GSW | None | No |
| 3 | CONTROL | NORMAL | 46 | Black | Male | 9 | 7.1 | Multiple injuries | None | No |
| 4 | CONTROL | NORMAL | 33 | White | Male | 15 | 7 | GSW | None | No |
| 5 | CONTROL | NORMAL | 48 | White | Male | 26 | 6.9 | ASCVD | None | No |
| 6 | CONTROL | NORMAL | 38 | Black | Male | 16 | 6.9 | Lung sarcoidosis | None | No |
| 7 | CONTROL | NORMAL | 65 | Black | Felame | 23 | 6.9 | ASCVD | None | No |
| 8 | CONTROL | NORMAL | 52 | White | Male | 30 | 7.3 | ASCVD | None | No |
| 9 | CONTROL | NORMAL | 63 | White | Female | 30 | 7.1 | Ovarian cancer | None | No |
| 10 | CONTROL | NORMAL | 37 | White | Male | 24 | 7 | ASCVD | None | No |
| 11 | CONTROL | NORMAL | 72 | White | Female | 23 | 6.9 | MVA | None | No |
| 12 | CONTROL | NORMAL | 42 | White | Female | 23 | 6.9 | Mitral valve prolapse | None | No |
| 13 | CONTROL | NORMAL | 31 | White | Male | 16 | 7.2 | MVA | None | No |
| 14 | CONTROL | NORMAL | 28 | White | Male | 13 | 6.8 | Electrocution | None | No |
| 15 | CONTROL | NORMAL | 53 | White | Male | 15 | 6.9 | ASCVD |  |  |
| 1 | SUICIDE | Major depression, alc | 27 | White | Male | 24 | 7 | GSW | None | NO |
| 2 | SUICIDE | Major depression, alc | 44 | White | Female | 11 | 7.2 | Drug overdose | Nortriptyline | YES |
| 3 | SUICIDE | Major depression | 24 | White | Male | 7 | 7.1 | GSW | Ethanol | NO |
| 4 | SUICIDE | Major Depression, Pc | 43 | White | Male | 12 | 7 | Drug overdose | Propoxyphene, Acetaminophen | NO |
| 5 | SUICIDE | Major depression | 53 | White | Male | 23 | 6.9 | Jumped 3rd floor | None | NO |
| 6 | SUICIDE | Major depression, Alc | 41 | White | Female | 27 | 7.1 | Drug overdose | Amitriptyline, Desipramine, Diphenhydramine, Nortriptyline | YES |
| 7 | SUICIDE | Major depression | 36 | White | Female | 18 | 7.2 | GSW | None | NO |
| 8 | SUICIDE | Major depression, Alc | 38 | White | Male | 24 | 7 | Drug overdose, Ethanol overdose | Ethanol, Diphenhydramine | NO |
| 9 | SUICIDE | Major depression (29 | 46 | White | Female | 16 | 6.8 | Drug overdose, Nortriptyline intoxication | Nortriptyline | YES |
| 10 | SUICIDE | Major depression (29 | 30 | White | Male | 17 | 7.1 | Hanging suicide | Effexor | YES |
| 11 | SUICIDE | Major depression (29 | 74 | White | Female | 27 | 7 | Suicide by Effexor OD | Effexor, Ethanol | YES |
| 12 | SUICIDE | Major depression (29 | 25 | White | Male | 14 | 6.8 | Suicide by hanging, asphyxia | Ethanol | NO |
| 13 | SUICIDE | Major depression, NC | 23 | Black | Male | 23 | 6.9 | Hanging suicide | None | NO |
| 14 | SUICIDE | Major depression (29 | 67 | White | Male | 22 | 7 | GSW to chest | Prozac, Effexor | YES |
| 15 | SUICIDE | Major depression (29 | 40 | White | Female | 20 | 7 | Suicide by OD | Acetaminophen, Hydrocodone, Diphenhydramine, Xanax | NO |
| 1' | Non-Suicide | Major depression, rec | 65 | White | Male | 14 | 6.9 | ASCVD | None | NO |
| 2' | Non-Suicide | Major depression, rec | 55 | Black | Female | 8 | 6.4 | ASCVD | Fluoxetine, Ethanol | YES |
| 3' | Non-Suicide | Major depression, rec | 71 | White | Male | 4 | 6.3 | ASCVD | Bupropion, Diltiazem | YES |
| 4' | Non-Suicide | Depression, NOS (31 | 74 | Black | Female | 7 | 6.7 | ASCVD | Paroxetine, Thioridazine | YES |
| 5' | Non-Suicide | Major depression, sin | 14 | White | Male | 11 | 7 | MVA | Sertraline | YES |
| 6' | Non-Suicide | Major depression, rec | 39 | White | Male | 36 | 6.8 | Fatty liver | Thioridazine | NO |
| 7' | Non-Suicide | Major depression, rec | 46 | Black | Male | 20 | 7.1 | Seizure disorder | Fluoxetine, Risperidone | YES |
| 8' | Non-Suicide | Major depression, rec | 59 | White | Male | 20 | 7 | ASCVD | Sertraline, Atropine | YES |
| 9' | Non-Suicide | Major depression, rec | 46 | White | Female | 23 | 6.9 | Mixed drug intoxication | Bupropion, Lamotrigine, Diphenhydramine | YES |
| 10' | Non-Suicide | Major depression, rec | 29 | White | Female | 22 | 6.9 | Morbid obesity, Cardiomegaly | Fluoxetine, Norfluoxetine, Norpropoxyphene | YES |
| 11' | Non-Suicide | Major depression, NC | 49 | White | Male | 24 | 7.1 | ASCVD | Desmethylsertraline | YES |
| 12' | Non-Suicide | Major depression, rec | 47 | White | Female | 26 | 6.5 | DKA | Fluoxetine | YES |

ASCVD, atherosclerotic cardiovascular disease; GSW, gunshot wound; MDD, major depressive disorder; MVA, motor vehicle accident Mean +/- SD age is 45.07 +/- 13.59 years; PMI is 18.40 +/- 7.84 hours; brain pH is 7.01 +/- 0.15; 5 Black, 10 White; 11 Males, 4 Females Mean +/- SD age is 40.73 +/- 15.04 years; PMI is 19.00 +/- 6.07 hours; brain pH is 7.01 +/- 0.12; 1 Black, 14 White; 9 Males, 6 Females Mean +/- SD age is 58.58 +/- 29.99 years; PMI is 17.91 +/- 16 hours; brain pH is 6.8 +/- 0.13; 3 Black, 9 White; 7 Males, 5 Females

All tissues from normal controls, depressed suicides and non-suicide subjects were screened for evidence of neuropathology. In addition, in each case, screening for the presence of HIV was done in blood samples, and all HIV-positive cases were excluded. Toxicology data were obtained by the analysis of urine and blood samples. pH of the brain was measured in cerebellum in all cases as described (Harrison et al., 1995). Psychiatric drugs in common use as well as drugs of abuse were screened for by using mass spectroscopy. Prescribed drugs were also screened for in interviews.

Control subjects with a known psychiatric illness or a history of alcohol or another drug abuse were excluded. However, alcohol or other substance abuse was present in the MDD subjects as indicated.

### Diagnostic Method

Families were queried on all medications or drugs of abuse by trained interviewers. At least one family member, after giving written informed consent, underwent an interview based on the Diagnostic Evaluation After Death (DEAD) (Zalcman and Endicott, 1983 and the Structured Clinical Interview for the DSM-IV (SCID) (Spitzer et al., 1992). This was done as described in a previous study (Donati et al., 2008).

### Sequential Detergent Extraction of Brain Membranes

Brain samples were dissected from the fresh brain and stored at −80°C or dissected from frozen brain tissue with a Stryker autopsy saw, repackaged, and stored at −80°C until use. Brain samples (Pre-Frontal Cortex-PFC) were resuspended and minced in TME buffer (10 mm Tris-HCl, 1 mm MgCl2, 1 mm EDTA, pH 7.5; ∼1 ml/100 mg tissue) followed by homogenization in a motorized Teflon glass homogenizer. Small amount of whole tissue homogenate (H) was saved to be run on western blot along with other cell fractions. The rest of the H samples were centrifuged at 100,000 × *g* for 1 h at 4°C, and the supernatant (cytosol) and pellet (Plasma membrane-PM) were saved.

The crude membrane pellet was extracted with 0.75 ml of TME containing 1% Triton X-100 for 1h at 4°C followed by homogenization as above. This sample was centrifuged as above and both the supernatant (TX-100 extract) and pellet (TX-100-resistant membrane fraction) were saved. This pellet was extracted with 0.75 ml of TME containing 1% Triton X-114 for 1h at 4°C and homogenized as above. The sample was centrifuged as above and both the supernatant (TX-114 extract) and pellet (detergent-insoluble pellet) were saved. The detergent-insoluble pellet could not be efficiently solubilized to be quantified. From here on out, the TX-100 extract will be referred to as the TX-100-soluble domain and the TX-114 extract will be referred to as the TX-100-resistant domain. All fractions were assayed for protein content (Bio-Rad Protein Assay; Bio-Rad, Hercules, CA) and frozen at −80°C until further use. Frontal cortex was the only brain region available for these experiments (Donati et al., 2008).

### SDS-PAGE and Western Blotting

Whole tissue homogenate (H), plasma membrane (PM), TX-100- and TX-114-soluble (TX-100-resistant) membrane fractions (12–15μg) were analyzed by SDS-PAGE followed by Western blotting. The gels were transferred to Nitrocellulose membranes (Bio-Rad, Hercules, CA USA) by

Western blotting. The membranes were blocked with 5% nonfat dry milk diluted in TBS-T (10 mM Tris–HCl, 159 mM NaCl, and 0.1% Tween 20, pH 7.4) for 1 h. Following the blocking step, membranes were washed with Tris-buffered saline/Tween 20 and then incubated with an anti-acetyl-α-tubulin (SIGMA-ALDRICH #T7451 Clone 6-11B-1), α-tubulin (SIGMA-ALDRICH #T9026), HDAC6 (Cell Signaling #7558S), ATAT-1 (SIGMA-ALDRICH #HPA046816), GAPDH (Proteintech #60004-1-Ig) overnight at 4°C. Membranes were washed with TBS-T and incubated with a secondary antibody [HRP-linked anti-mouse antibody IgG F(ab′)2 or HRP-linked anti-rabbit antibody IgG F(ab′)2 (Jackson ImmunoResearch, West Grove, PA, USA, catalog #115-036-072 for mouse, and catalog #111-036-047 for rabbit, RRID) for 1 h at room temperature, washed, and developed using ECL Luminata Forte chemiluminescent reagent (Millipore, Billerica, MA, USA). Blots were imaged using Chemidoc computerized densitometer (Bio-Rad, Hercules, CA, USA). The signal intensity of bands from each image were quantitated by densitometry using Image-J software (NIH) and the TX-100-resistant acetyl-α-tubulin/α-tubulin (TX-114) was compared. The acetyl-α-tubulin/α-tubulin were also observed in plasma membrane (PM) from Control (NC), depressed Suicide (DS) and depressed non-suicide (DNS) samples as described (Toki et al., 1999; Donati et al., 2008). Additionally, HDAC6, ATAT-1 and GAPDH expression differences were analyzed between the 3 groups (C, DS & DNS).

### Normalization

To be consistent throughout the data collection, same amount of starting material (H) was used for membrane isolation and lipid-raft extraction. Additionally, GAPDH was used as loading control for all 3 groups to account for expression differences in α-tubulin, HDAC6 and ATAT-1 amongst groups. Additionally, this normalization procedure was repeated when comparing the amount of acetyl-α-tubulin/α-tubulin (normalized densitometry value = sample value/mean value). This allowed us to compare samples accurately among gels and their corresponding blots.

### Statistical Methods

Western blot data were analyzed for statistical significance by unpaired, two-tailed Student’s *t* test or one-way ANOVA using Prism 4.0 software package for statistical data analysis (Graph Pad, San Diego, CA). Means are ±SEM, and differences for all experiments were considered significant at *p* < 0.05 (\**p* < 0.05; \*\**p* < 0.02). The differences in TX-114 acetyl-α-tubulin/α-tubulin, age, gender, pH of the brain, and postmortem interval (PMI) between depressed and control subjects were analyzed using the independent-sample *t* test. The relationships between TX-114 acetyl-α-tubulin/α-tubulin and PMI, and age were determined by Pearson product-moment correlation analysis. Values of *p* were two-tailed. During data analysis, confounding variables such as age, PMI, gender, race and pH of the brain were also used as covariates (Proc GLM)(SAS 9.4 statistical software package). A linear model was used to compare NC, DS, and DNS subjects simultaneously adjusting the effects of age, gender, postmortem interval (PMI), brain pH, antidepressant use, ethanol use, non-psychotropic medicine use, violent-suicide and Hypoxia. For post-hoc multiple comparisons, we used Bonferroni (Dunn) t Tests to adjust the type I error rates, and we reported mean differences (mean-diff) and confidence interval (CI) to test the significance at the 0.05 level. In addition, each outcome measure was tested for normality (Kolmogorov-Smirnov) before running the model. All results are included in tables 2 and 3. Table 2 shows the overall model

**Table 2:** ANCOVA and multiple comparisons of results

**ANCOVA**
| **Dependent Variable** | **DF (Model, Error)** | **F Value** | **Pr > F** |
| --- | --- | --- | --- |
| **acetyl-α tubulin/total $\alpha$ - tubulin (Homogenate)** | 8,32 | .89 | 0.57 |
| **acetyl-α tubulin/total $\alpha$ - tubulin (Plasma membrane)** | 8,32 | 2.17 | 0.04 |
| **acetyl-α tubulin/total $\alpha$ - tubulin (Lipid-rafts)** | 8,32 | 6.51 | <.0001 |

| **DV** | **Group Comparison** | **Difference Between Means** | **Simultaneous 95% Confidence Limits** |  |  |
| --- | --- | --- | --- | --- | --- |
| **acetyl-α tubulin/total $\alpha$ - tubulin (Homogenate)** | NC - DS | 0.029 | -0.7510 | 0.8091 |  |
|  | NC - DNS | 0.6143 | -0.2115 | 1.44 |  |
|  | DS - DNS | 0.5853 | -0.2277 | 1.3982 |  |
| **acetyl-α tubulin/total $\alpha$ - tubulin (Plasma Membrane)** | NC - DNS | 0.5592 | 0.1544 | 0.9640 | *** |
|  | NC - DS | 0.5947 | 0.2124 | 0.9771 | *** |
|  | DNS - DS | 0.0355 | -0.3630 | 0.4340 |  |
| **acetyl-α tubulin/total $\alpha$ - tubulin (Lipid-rafts)** | NC - DS | 3.9386 | 2.4972 | 5.3800 | *** |
|  | NC - DNS | 4.0179 | 2.4920 | 5.5439 | *** |
|  | DS - DNS | 0.0793 | -1.5816 | 1.4230 |  |

## Results

There were 11 males and 4 females in the NC group, 9 males and 6 females in the DS group and 7 males and 5 females in the DNS groups (Table 1). The age range was 14-74 years, whereas the postmortem interval (PMI) was in the range of 5-30 h. There were no significant differences in age (t=.83; df =26; p =.29) or PMI (t=-.23; df = 28; p=.82) between suicides and normal control subjects. The mean brain pH values of NC, DS and DNS were 7.01± .14, 7.01± .12 and 6.8±.13 respectively, which were not different (t=.14; df =28; p=.89).

### Prefrontal cortex postmortem tissue from control, depressed suicide and depressed non-suicide subjects showed no changes in acetylation of α-tubulin in whole tissue homogenate

The whole tissue homogenate (H) sample derived before plasma membrane and lipid-raft isolation from prefrontal cortex tissue of control (n=15), depressed suicides (n=15) and depressed non-suicides (n=12) showed no changes in acetylated-α-tubulin (Figure 1A, B, C & D). The quantification of the results from all three groups NC, DS & DNS showed no significant differences the extent of tubulin acetylation or any significant significant effects of covariates (table 2).

**Figure 1:**
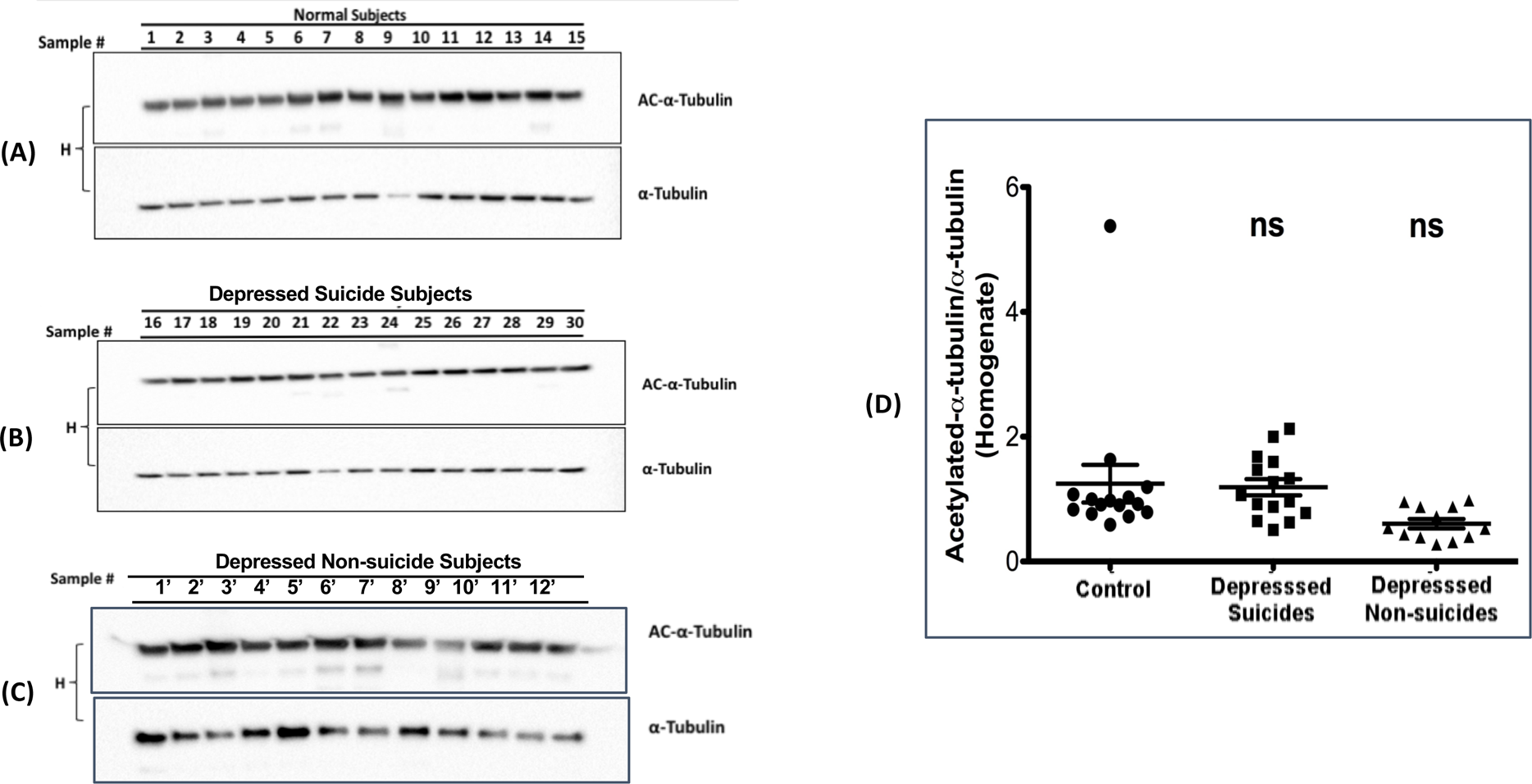
**α-tubulin acetylation status in postmortem brain prefrontal cortex derived from normal and depressed suicides**: Prefrontal cortex tissue from control (normal subjects), depressed suicides and depressed non-suicides were homogenized (H), run on SDS-PAGE gel and transferred to nitrocellulose for detection with either acetyl-α-tubulin or α-tubulin antibodies. The the signal intensity was quantified and scatter plots are used to show the extent of tubulin acetylation in each group (ns=non-significant compared to control).

### Depressed suicide brain plasma membrane localized tubulin shows decreased acetylation of α-tubulin compared to that of in normal controls

Plasma membranes isolated from prefrontal cortex postmortem tissue of NC, DS & DNS were compared for acetylation status of membrane-associated tubulin. Five samples from each group (NC & DS) were loaded on a single gel (Figure 2A, B, & C). Additionally, DNS samples (protein concentration equal to NC & DS group subjects) were loaded on a separate gel (Figure 2D). SDS-PAGE analysis showed significant decrease in acetyl-α-tubulin in DS subjects (1-15) and DNS subjects (n=12) compared to the NC subjects. Significant changes were observed between groups in acetyl-α-tubulin/α-tubulin (F(2) =8.79, p=.0009). The tests from multiple comparisons showed significant differences at 95% confidence level between control vs DS (mean-diff = 0.59, CI = (0.21, 0.97)) and NC vs DNS (mean-diff = 0.56, CI = (.15, 0.96)) (Figure 1E, table3). There were no significant effects of any covariates.

**Figure 2:**
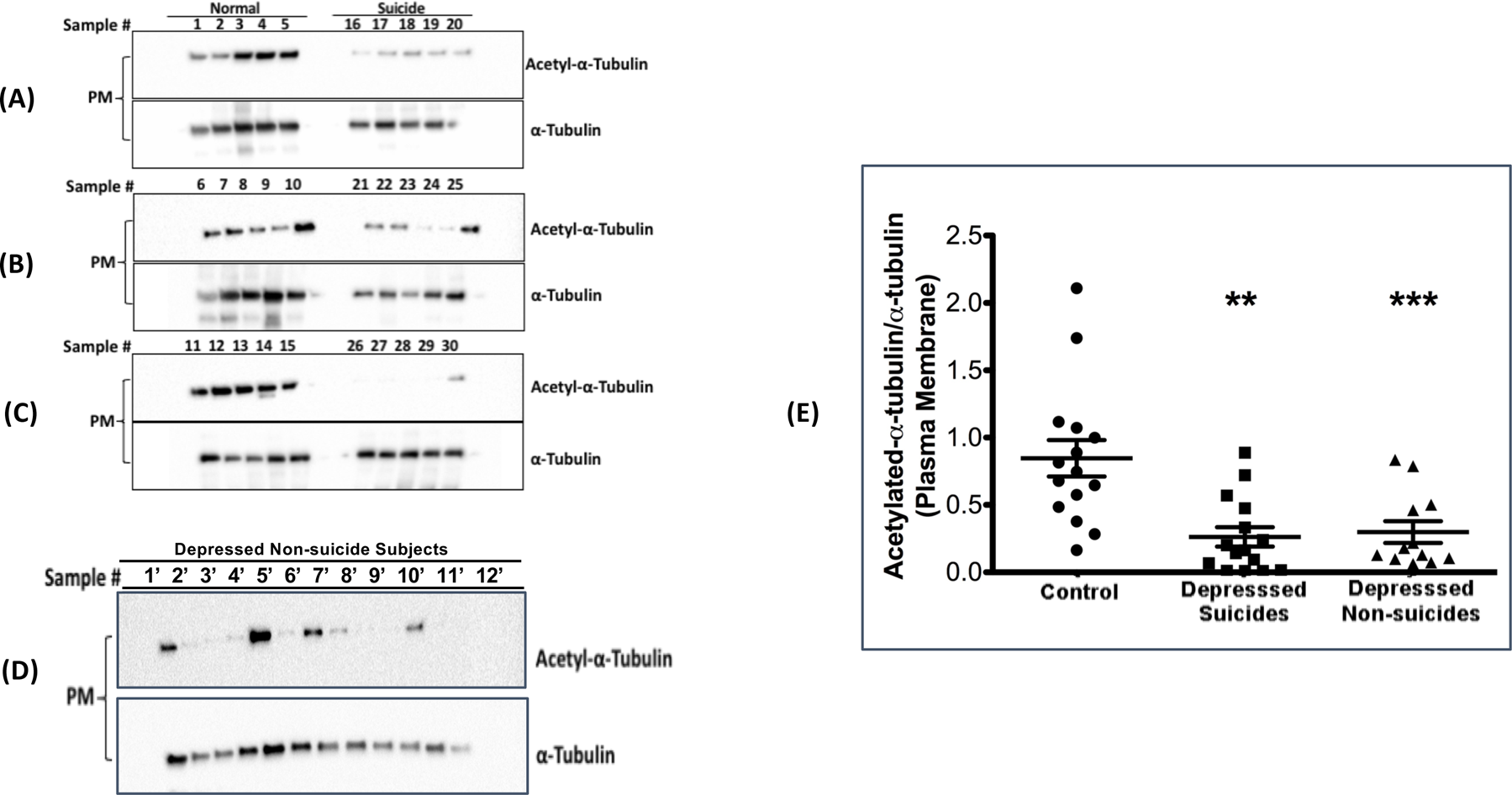
**Acetylated tubulin in plasma membrane prepared from prefrontal cortex is decreased in suicides relative to control**: Plasma membrane (PM) was isolated from the samples presented in figure 1 and analyzed in the same manner. Scatter plots are used to show the spread of tubulin modification in both the groups (*** p=.0001).

### Detergent-resistant/lipid-raft membrane domains as well as TritonX-114-resistant/non-raft domains show decreased acetylation of α-tubulin in depressed subjects compared to normal control postmortem prefrontal cortex

Using plasma membrane as the starting material (Figure 2) we isolated lipid-raft fractions in order to determine whether the decrease in acetylated tubulin was localized to lipid-rafts (Figure 3A & B). The raft domains showed differences in levels of tubulin acetylation. The quantification of the results from all three groups Control, DS & DNS showed significant differences between acetyl-α-tubulin/α-tubulin levels in Detergent-resistant lipid-rafts (F(2) = 6.51, P<.0001) (Figure 3C). The multiple comparisons between control vs depressed suicide subjects and control vs depressed non-suicides showed significant differences between the extent of acetyl-α-tubulin/α-tubulin in detergent-resistant lipid-rafts [Control vs DS (mean-diff = 3.94, CI = (2.49,5.38)), Control vs DNS (mean-diff = 4.02, CI = (2.49, 5.54))] as shown in table3. There was a significant effect of hypoxia on lipid-raft tubulin (t=-2.95, p=.01)

**Figure 3:**
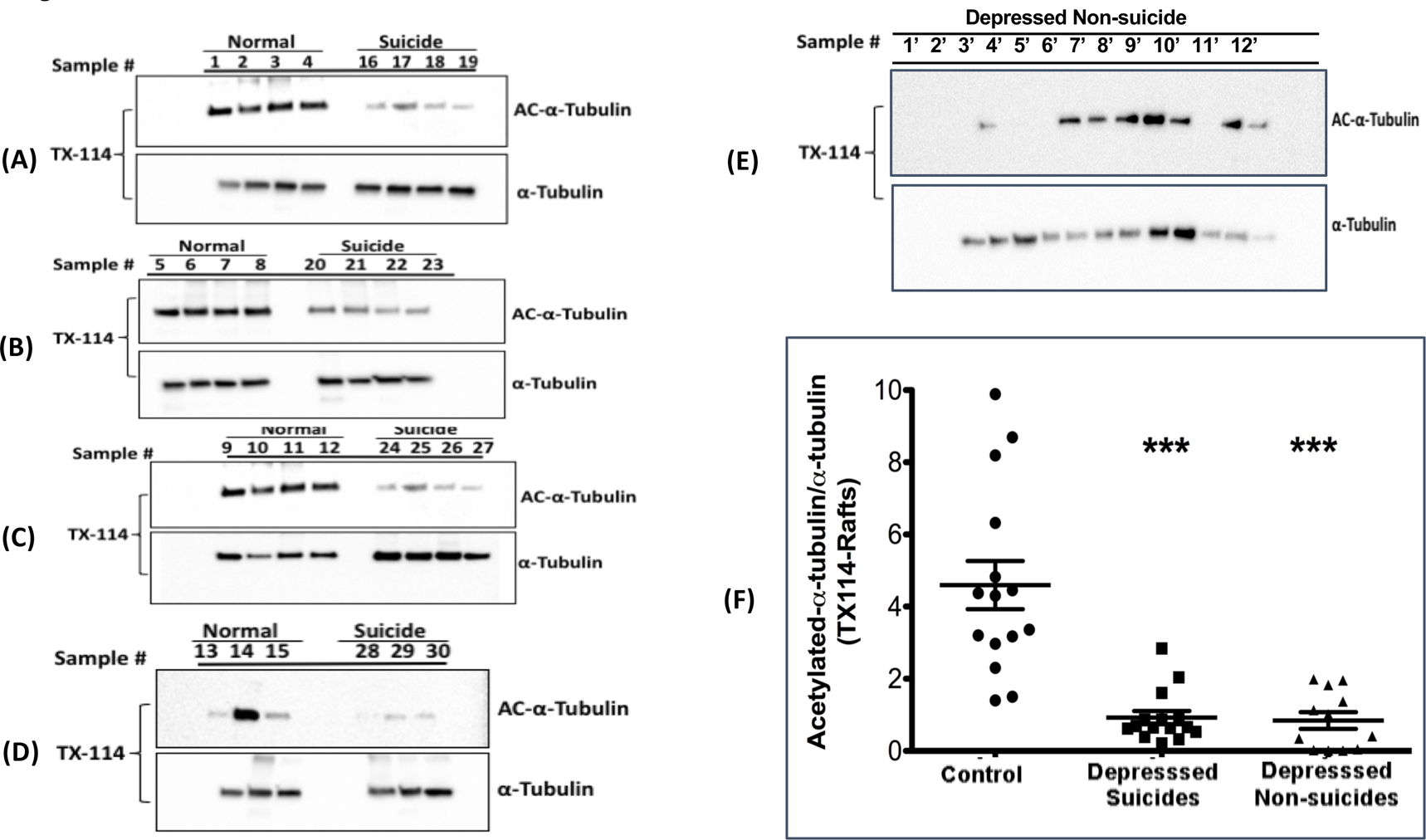
**Acetylated tubulin in lipid-rafts prepared from prefrontal cortex plasma membrane is decreased in suicides relative to control**: Plasma membranes were purified and lipid rafts were prepared by TritonX-100-resistant (lipid-rafts) and TritonX-114-resistant (non-rafts) micro-domain isolation. Samples were analyzed as in figures 1 and 2. Tubulin and acetylated tubulin were quantified and scatter plots were used to show the distribution of tubulin modification in both the groups (C) (*** P<0.0001)

### Neither tubulin acetylating nor tubulin deacetylating anzymes show altered expression in depressed brain

HDAC6 regulates deacetylation of α-tubulin and previous studies in blood cells and postmortem brain tissue derived from patients with mood disorders showed altered HDAC6 expression (Covington et al., 2009). We did not observe these changes. (Figure 4A, B, C, D). The enzyme ATAT-1 specifically acetylates α-tubulin at K-40, whereas HDAC6 deacetylates. Therefore, along with studying changes in HDAC6 expression levels, we investigated ATAT-1 enzyme level changes. ATAT-1 expression levels/GAPDH remain statistically non-significant amongst NC, DNS & DS (F(2) = .96, P=.39 (Figure 4 A,B,C, E). We investigated further the effect of GAPDH or any other co-variates on HDAC6 and found no significant effect (Figure 4D). Similarly, we investigated whether GAPDH and other covariates have any effect on ATAT-1. For one unit increase in GAPDH, ATAT-1 is increasing by .20 unit but not significantly (t=.47, p=.64). Hypoxia (t= −2.25, p=.03) and Violent Suicide (t=2.44, t=.02) have a significant effect on ATAT1. However, there are no group differences in the overall model (F(2)=1.87, p=.17) Most importantly, the ATAT1/HDAC6 ratio is not significantly different amongst the three groups (figure 4F), suggesting that there is no meaningful change in the expression of the enzymes regulating tubulin acetylation.

**Figure 4:**
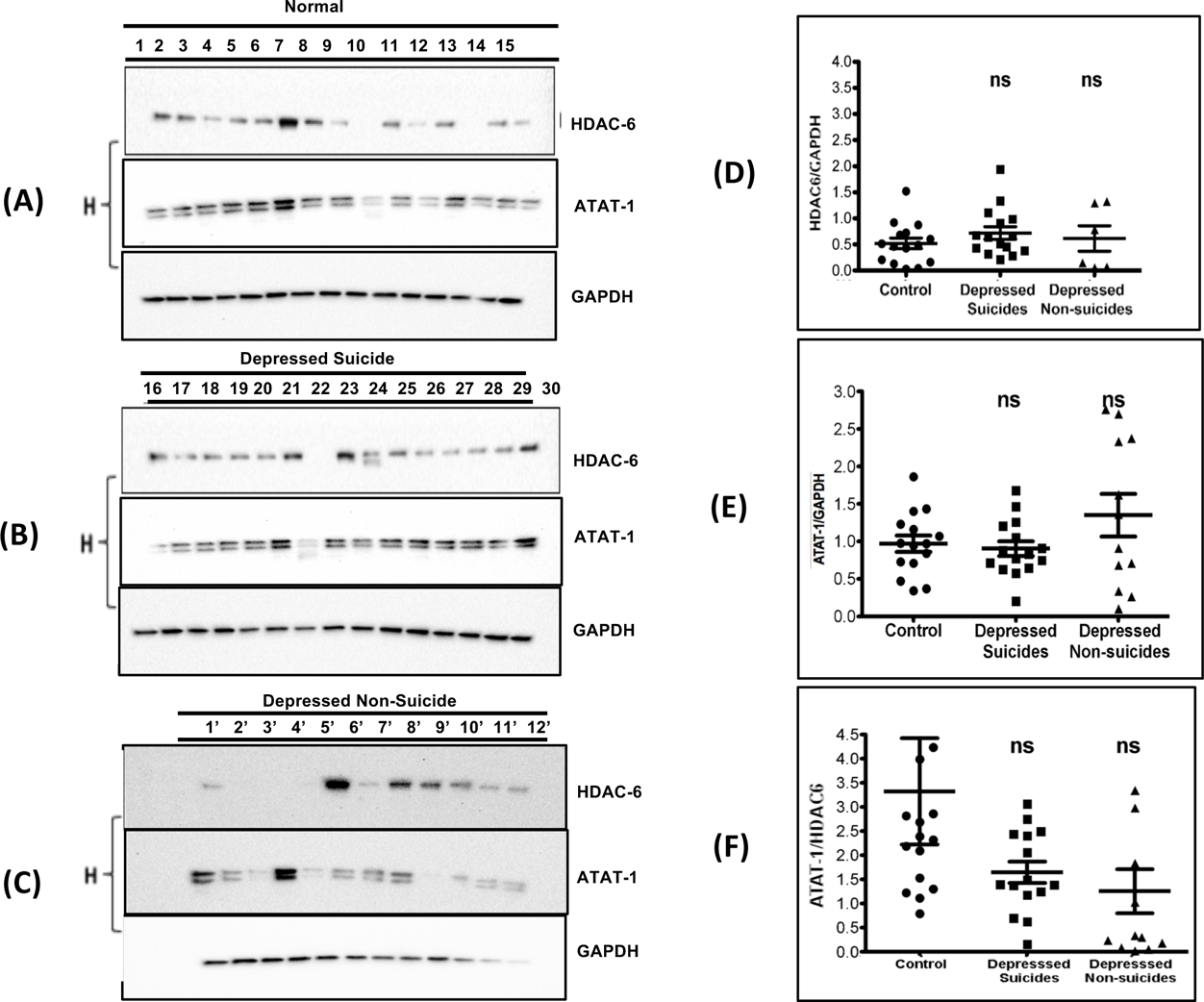
**Expression of tubulin acetylating and deacetylating enzymes in postmortem tissue**: Tissue homogenates were analyzed for presence of ATAT-1 (acetylating) and HDAC-6 (deacetyating) enzymes in postmortem homogenates (as in figure 1). Ratios of each pair were calculated and plotted in G & H.

## Discussion

Postmortem results presented here dovetail well with results in a cellular model revealing that increased tubulin acetylation causes the antidepressant signature response of Gα_s_ translocation from lipid rafts (Singh et al., 2018) Current findings in post-mortem brain tissue suggest that acetylation status of tubulin may be important for sequestration of Gα_s_ in lipid rafts, as seen in depression (Donati et al). The findings lend a molecular rationale to antidepressant effects observed in HDAC6 depleted (Espallergues et al., 2012; Fukada et al., 2012; Lee et al., 2012) or pharmacological inhibitor treated animals (Jochems et al., 2014), where increased tubulin acetylation induced behavioral effects similar to that of traditional antidepressants.

Importantly, tubulin posttranslational modifications are observed in postmortem brain tissue from MDD subjects, resulting in abnormal cytoskeletal organization, disruption of microtubule dynamics, which are essential for neurite growth, synaptogenesis and dendritic arborization (Wong et al., 2013). Furthermore, proteomic studies from postmortem brain tissue of MDD subjects showed changes in proteins involved in cytoskeletal arrangement, neurotransmission and synaptic function (Scifo et al., 2017). Chronic stress results in dendritic retraction and synaptic density loss causing regional atrophy in the hippocampus, amygdala, and prefrontal cortex, as detected in MRI scans of psychiatric patients (McEwen et al., 2015). Finally, there is literature suggesting that microtubules might convey mood and consciousness (Cocchi et al., 2010). Based on these data, altered tubulin and microtubules appear to be a common parameter for several neuropsychiatric disorders. α-tubulin undergoes acetylation and deacetylation at Lysine-40 (K40), catalyzed by acetyl transferase and deacetylase enzymes respectively. Histone deacetylase-6 (HDAC6), a cytosolic HDAC is known to deacetylate 〈-tubulin. HDAC6 enzyme is highly expressed in brain, where it is known to regulate emotional behaviors in rodents. HDAC6-deficient mice display hyperactivity, low anxiety, and low depressive like phenotype indicating that acetylation status maintains the cellular activity associated with control of emotions (Fukada et al., 2012). Similarly, pharmacological inhibition of HDAC6 in rodents using inhibitors with increased brain bioavailability (ACY738, ACY-775) show increased anxiolytic and antidepressant-like effects in mice undergoing “depression–inducing” paradigms (Jochems et al., 2014). Furthermore, chronic stress in rodents has been shown to induce increased expression of HDAC6 in hippocampus (Jianhua et al., 2017). Decreased levels of acetylated tubulin are found in the hippocampus of rats following social isolation (Bianchi et al., 2009). These studies further corroborated the microtubule roles, especially tubulin acetylation, in the pathophysiology of depression. Decreased dendritic spine density and reduced dendritic arborization are associated with neurological diseases (Blanpied and Ehlers, 2004; Penzes and Vanleeuwen, 2011), including intellectual disability (Kaufmann et al., 2000), depression (Duman and Canli, 2015) and schizophrenia (Penzes and Vanleeuwen, 2011; Glausier and Lewis, 2013). Chronic stress induces atrophy in hippocampus and prefrontal cortex, areas important for mood regulation. Reduced dendritic field size results in abrogated synaptogenesis (Gold, 2015). HDAC6 regulates deacetylation of α-tubulin and previous studies in blood cells and postmortem brain tissue derived from patients with mood disorders showed altered HDAC6 expression (Covington et al., 2009). Post-translational modifications in α-tubulin (acetyl-α-tubulin) result from either increased enzyme expression or increased enzyme activity. We did not observe any specific expression pattern within each group or amongst three groups when normalized to total α-tubulin (Control, DS, DNS). The enzyme ATAT-1 specifically acetylates the α-tubulin at K-40, acting as the “accelerator” to the “brake” represented by HDAC6. ATAT-1 expression levels show no significant difference amongst control, depressed suicides and depressed non-suicides (F(8,32) = 1.04, P=.43) (Figure 4). Nonetheless, results in figures 2 and 3 reveal that depressed subjects show decreased acetylated a tubulin in membrane fractions. This suggests that the activity of HDAC6 relative to ATAT1 is increased without any change in the expression of either enzyme. This could be explained by multiple factors. First, HDAC6 is regulated by nitrosylation (Okuda et.al, 2015). Perhaps more importantly, only membrane tubulin (particularly lipid-raft tubulin) was affected, as the total degree of tubulin acetylation was constant amongst all groups. Perhaps some membrane translocating mechanism is at play.

These findings are consistent with a link between decreased α-tubulin acetylation and increased localization of Gα_s_ in lipid-rafts. Our *in vitro* studies in C6 cells showing HDAC6 inhibition induced α-tubulin acetylation results in disruption of tubulin-Gα_s_ complex, specifically in the lipid-raft domain, bolster this (Singh et al., 2018). Furthermore, membrane tubulin appears to be associated, preferentially, with lipid rafts (Goudenege et.al, 2007), so “membrane tubulin and lipid-raft tubulin may be identical. While earlier studies showed that tubulin binding to Gα_s_ was sensitive to Gα_s_ conformation, the nucleotide status of tubulin was not important (Yu et al., 1999). The apparent binding site for Gα_s_ on tubulin involves the α3β5 region of Gα_s_ and the GTP-binding pocket of ®-tubulin (Layden et al., 2008; Dave et al., 2011) While the structural changes to ®-tubulin resulting from modifying 〈-tubulin have not been established, it is clear that modifying 〈-tubulin has structural implications for the dimer (Nogales et al., 1998).

This study strikes a thematic note in revealing that compounds with antidepressant activity show a consistent “biosignature” in the release of Gα_s_ from lipid rafts and the subsequent association of that molecule with adenylyl cyclase, evoking a sustained increase in cellular cAMP (Singh et al., 2018). We have also demonstrated that increased acetylation of tubulin can explain this, in part. Furthermore, the diminished tubulin acetylation seen in lipid rafts from depressed subjects might explain the increase in Gα_s_ seen in their lipid rafts. Nevertheless, the ability of monoamine-centered antidepressants to mitigate Gα_s_-tubulin association without altering tubulin acetylation (Singh et al., 2018) argues for the complexity of depression and its therapy.

## Acknowledgements

Support was provided by VA Merit award-BX001149 (MMR); NIH RO1AT009169 (MMR);. NIH R21 NS 109862 (MMR); RO1MH106565 (GNP). American Heart Association (AHA) Postdoctoral award-16POST27770113 (Harinder Singh). MMR is a VA Research Career Scientist. The authors thank Miljiana Petkovic for technical expertise.

